# MousePZT: A Simple, Reliable, Low-Cost Device for Vital Sign Monitoring and Respiratory Gating in Mice Under Anesthesia

**DOI:** 10.1101/2023.02.23.529566

**Authors:** Daniel A. Rivera, Anne E. Buglione, Chris B. Schaffer

## Abstract

Small animal studies in biomedical research often require anesthesia to reduce pain or stress experienced by research animals and to minimize motion artifact for imaging or other measurements. Anesthetized animals should be closely monitored to avoid complications and unintended effects of altered physiology during such procedures. Many currently available monitoring devices can be expensive, invasive, or interfere with experimental design. Here, we present a low-cost device, based on a simple piezoelectric sensor, with a custom circuit and computer software that allows for measurements of both respiratory rate and heart rate in a non-invasive, minimal contact manner. We find the accuracy of the MousePZT device in measuring respiratory and heart rate under anesthesia and with pharmacologically induced changes in heart rate match those of commercial or more invasive systems. We also demonstrate that changes in respiratory rate can precede changes in heart rate associated with alterations in anesthetic depth. Additional circuitry on the device outputs a respiration locked trigger signal for respiratory-gating of imaging or other data acquisition that has high sensitivity and specificity for detecting respiratory cycles. We provide detailed construction documents and all necessary microcontroller and computer software, enabling straightforward adoption of this device.

## Introduction

The use of anesthetics in mice and other small laboratory animals is essential for many experimental procedures, but comes with risks, even in healthy animals. Isoflurane is an inhalant anesthetic commonly used due to its rapid-onset, short recovery time, and the ease of adjusting the anesthetic depth (*1, 2*). The effective anesthetic dose, however, depends on a variety of animal-dependent factors, so that fixing the percentage of isoflurane in the inhaled gas mixture can lead to over- and under-dosing of some animals. Insufficient or ‘light’ anesthesia risks awareness during stressful or painful procedures, while excessive or ‘deep’ anesthesia can lead to dangerous decreases in heart rate, respiratory rate, and blood pressure (*1, 2*). In addition, anesthetic agents alter a range of physiological parameters that can impact experimental results if not well-controlled across animals in an experiment. For example, in neurological and neurovascular research, isoflurane has been implicated in burst suppression (*3–5*), decreased cerebral metabolism (*3, 5, 6*), increased vasodilation (*3, 4, 6*), suppressed vasomotion (*6, 7*), altered neurovascular coupling responses (*3, 4, 6*), and neuroprotection against ischemia (*3*). Minimizing the variability in experiments due to these effects requires a narrowly restricted range of anesthetic depth. In addition, volatile anesthetics like isoflurane can accumulate in the body due to their fat solubility (*8*), necessitating lowered inhaled isoflurane doses over time to maintain the same level of anesthetic depth (*9*). Because vital signs, such as heart and respiratory rates, depend sensitively on anesthetic depth, monitoring these parameters and adjusting anesthetic dose to keep them steady is one approach to ensure appropriate anesthetic depth, detect side effects of administered drugs, and identify acute adverse events and trends during any procedure. In addition to monitoring vital signs, many live animal imaging modalities, such as MRI, micro-CT, PET, and optical or ultrasound imaging benefit from the ability to synchronize data acquisition with the respiratory cycle of the animal to minimize motion artifact, requiring a trigger signal indicating inspiration that can gate image acquisition (*10*).

Several tools and techniques are available to monitor vital signs, including pulse-oximeters, electrocardiogram (ECG), blood pressure cuffs, arterial catheters, capnographs, and thermometers (*11*). Labs may abstain from using some of these tools due to their invasiveness, the complexity of incorporating the tools into an experiment, poor reliability under anesthesia, or the overall cost. For example, arterial catheters provide excellent measures of blood pressure but are very invasive, requiring surgical implantation, and are typically suited only for terminal procedures. Rodent blood pressure cuffs placed around the tail measure blood pressure in the tail artery, but require warming the tail and often underestimate blood pressure, especially in anesthetized animals (*12*). ECG requires at least three leads be attached to the limbs by superficial clips, subcutaneous needles, or contact with a conductive platform. Although clips can be modified to reduce skin damage and conductive platforms are atraumatic, the additional wires and restricted positioning can quickly complicate the experimental setup, and the relatively weak ECG signal is prone to electrical interference. For pulse-oximetry, animals need to be prepped by removing any dark fur over the hindlimb or neck for clear optical access, signal amplitude depends sensitively on sensor placement, often requiring repeat adjustment, and the pressure from some sensors (e.g. thigh clips) can sometimes cause minor skin/soft tissue injury. While capnographs are simple to incorporate into exhalation lines, they produce more reliable signals with intubated or tracheotomized mice, as compared to free breathing animals, require additional calibration steps, and can be costly. The live animal research community would benefit from simple, reliable, and low-cost systems to monitor and record vital signs of mice that can be straightforwardly incorporated into surgical and experimental procedures.

Respiration is one of the simplest vital signs to measure and provides a sensitive measure of anesthetic depth (*11*). In many circumstances, respiratory rate can periodically be assessed visually by the experimenter. However, some experiments prevent continuous, direct visualization of the animal, and this approach may be error prone and forgoes a thorough log of respiratory rate through the experiment. Approaches to quantify respiration include electromyography to measure respiratory muscle activity, face masks or temperature probes to measure air movement, and cameras or movement sensors to detect body movement (*13*). Previously developed low-contact or low-cost respiratory monitoring systems for use in research utilized changes in infrared light reflectivity (*14*), amplitude and phase of reflected radio frequency signals (*15, 16*) due to chest wall motion, temperature due to expired air (*17*), and, more commonly, voltage signals from piezoelectric transducers (PZT) that were abutting the mouse (*18–23*). PZTs have the advantage of being highly sensitive to pressure perturbations due to breathing while also being low-cost. Few of the previously developed piezo-based sensors, however, are able to detect both respiratory rate and heart rate. Here, we present a simple, low-cost PZT-based system, MousePZT, that allows for measurement of both respiratory rate and heart rate with accuracy matching that of ECG and pulse-oximetry. The system has additional circuitry for real-time detection of respiration with high specificity and sensitivity, providing the respiratory gating capabilities so essential for many small animal imaging experiments.

## Results

The MousePZT system to measure mouse respiratory and heart rate uses a piezoelectric disk placed below the anesthetized mouse (Fig. 1A). The electrical signals produced by deflections of the chest or abdomen are amplified, filtered, and the voltage values are output by a microcontroller (Fig. 1B) to a computer app for signal processing to determine the respiratory and heart rate. MousePZT also includes an analog-based circuit (Fig. 1C) for respiratory peak detection that outputs a trigger signal (Fig. 1D), which would allow for data acquisition gating in a variety of applications.

**Figure 1.**
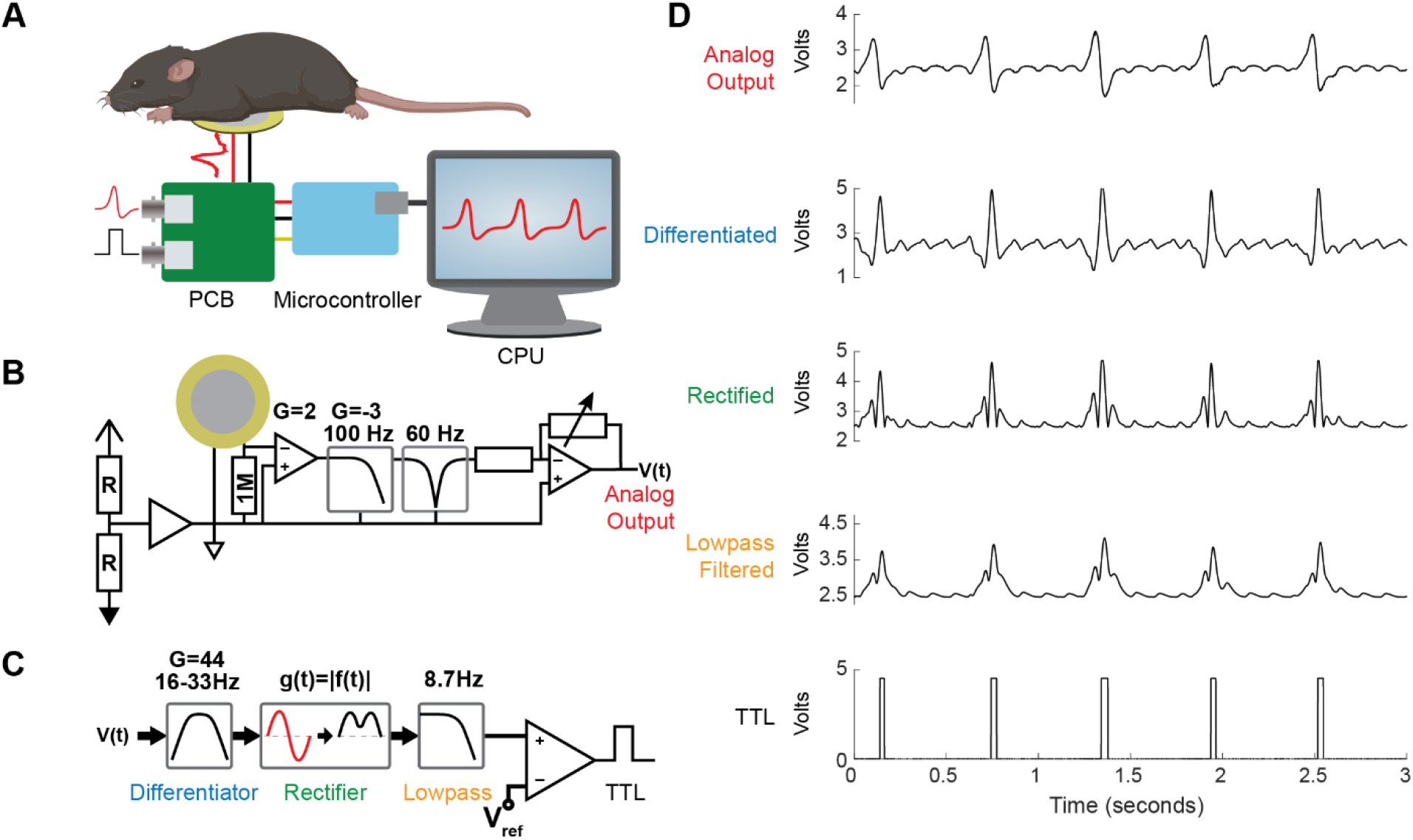
MousePZT vital sign monitoring system for mice. **(A)** Experimental setup for the system. The mouse rests on a piezoelectric sensor and the signal is amplified and filtered by a custom circuit, recorded with a microcontroller, and transmitted to a computer. **(B)** Simplified schematic for the custom circuit. The piezoelectric signal is amplified and lowpass filtered at 100 Hz, notch filtered to reduce line noise (50 or 60 Hz), and variably amplified. **(C)** Schematic of respiratory peak detection circuit. Signal from circuit in (B) is differentiated, rectified, and lowpass filtered before being passed to a comparator with a variable threshold. **(D)** Examples of a typical signal at each stage of the circuitry, with colored labels corresponding to circuit locations in (B) and (C).

### Testing the Accuracy of the MousePZT Respiratory Trigger

We tested the utility of the respiratory waveforms and the accuracy of the respiratory trigger circuit in mice anesthetized with isoflurane, an anesthetic widely used in laboratory settings. The signal was recorded with the mice positioned in dorsal, lateral, and ventral recumbency to demonstrate that the signal profile was recognizable and repeatable under various conditions (Fig. 2A). The output TTL signal from the respiratory trigger circuit matched well with respiratory epochs determined by digital signal processing of the analog circuit output (Fig. 2B). Session-to-session sensitivity and specificity were 98.6% and 96.0%, respectively, based on the presence or absence of TTL pulses during respiratory or inter-respiratory epochs. The average respiratory rate determined by output TTL pulses was highly correlated (R^2^ = 0.997) with that determined from the analog circuit output (Fig. 2C). We further compared the respiratory trigger with visually assessed respiration, based on excursions of the body wall. Two mice were recorded for 15-30 s in dorsal, lateral, and ventral recumbency to demonstrate the sensitivity of the PZTs regardless of the orientation of the animals. Based on visual scoring, there were no missed respirations, and no erroneous pulses detected (n=133 breaths, CI = [0 - 0.027]). Taken together, these data suggest high reliability and accuracy of the respiratory trigger circuit as well as the quantification of respiratory rate.

**Figure 2.**
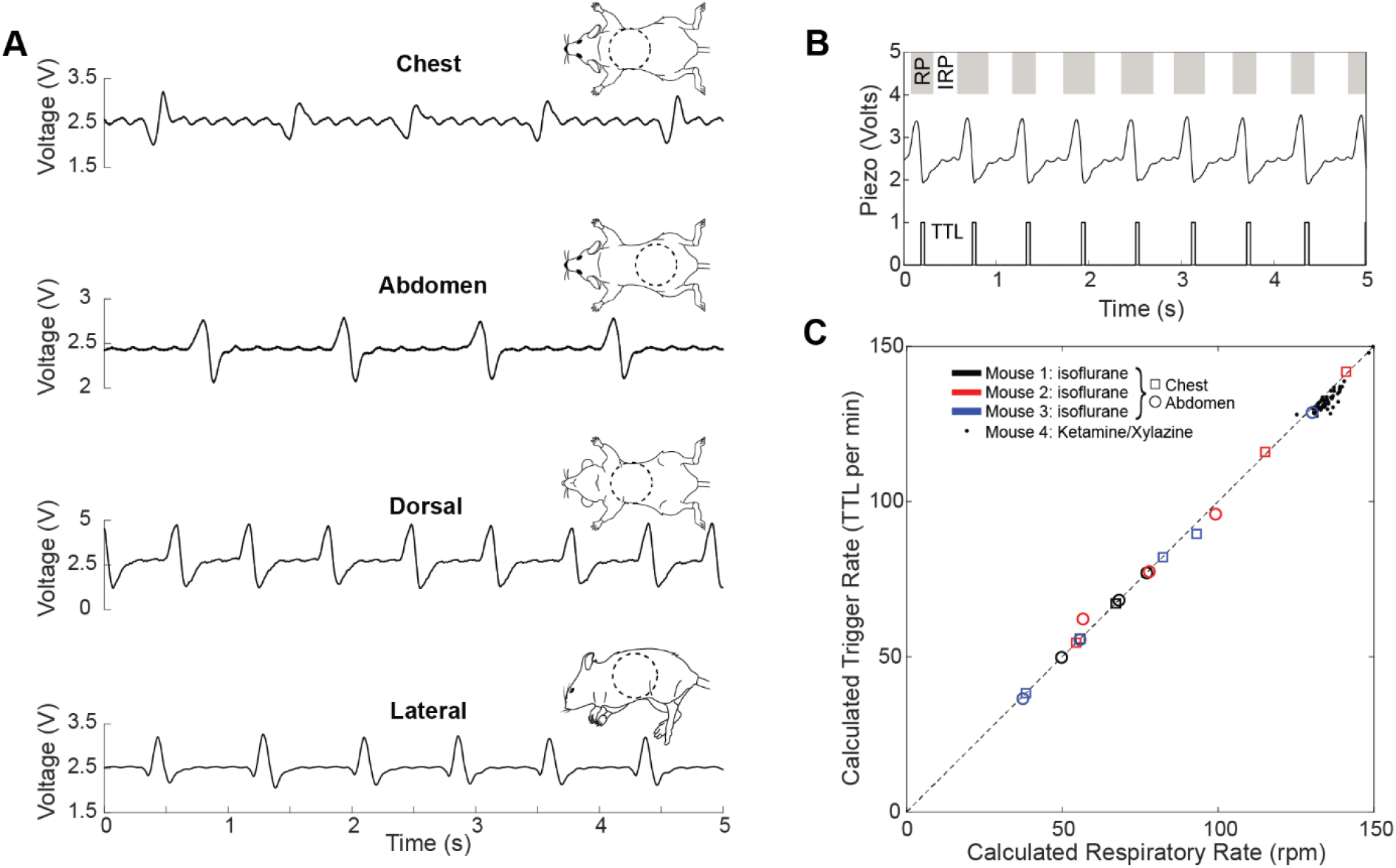
MousePZT provides a robust and accurate measurement of respiratory rate as well as a respiration locked trigger signal. **(A)** Examples of MousePZT signal with different sensor placement and animal orientation, as indicated in the schematics to the right of the traces (seen from above with sensor placement outlined by the dashed lines). **(B)** Example of the respiratory trigger. Respiratory phase (RP, gray) and inter-respiratory phases (IRP, white) are shown at top. Output piezo signal is shown in the middle, and the TTL respiratory trigger is at the bottom. **(C)** Correlation between breathing rates detected in 60-s intervals using the analog respiratory trigger circuit (Calculated Trigger Rate) and that from the MousePZT signal processed in MATLAB (Calculated Respiratory Rate) (R^2^ = 0.997). Plot includes data from three mice anesthetized on isoflurane with both chest and abdominal sensor placements, as well as 60 60-s epochs from a mouse anesthetized on ketamine/xylazine with a chest sensor placement.

### Confirmation of Heart Signal Presence in PZT with ECG

Within the PZT signal, there were noticeable smaller fluctuations between the respiratory peaks, which were even more prominent when the sensor was placed underneath the chest (Fig. 2A, *top*). We tested whether these smaller fluctuations originate from the contractions of the heart by simultaneously recording ECG. The small bumps in the PZT signal were locked to and occurred shortly after the QRS complex from the ECG signal (Fig. 3A, *top*). To calculate heart rate, we found the times of all heart beats and respirations, then the times of just the respirations, and finally calculated the median heart rate by measuring the interval between heart beats, while excluding data from times when the respiratory peak obscured heartbeat peaks (Fig. 3A, *bottom*; see Methods). With this approach, the absolute error between the PZT and the ECG based measurement of heart rate was only 1.4 ± 0.6 beats per minute (bpm), or a relative error of 0.24 ± 0.1%.

**Figure 3.**
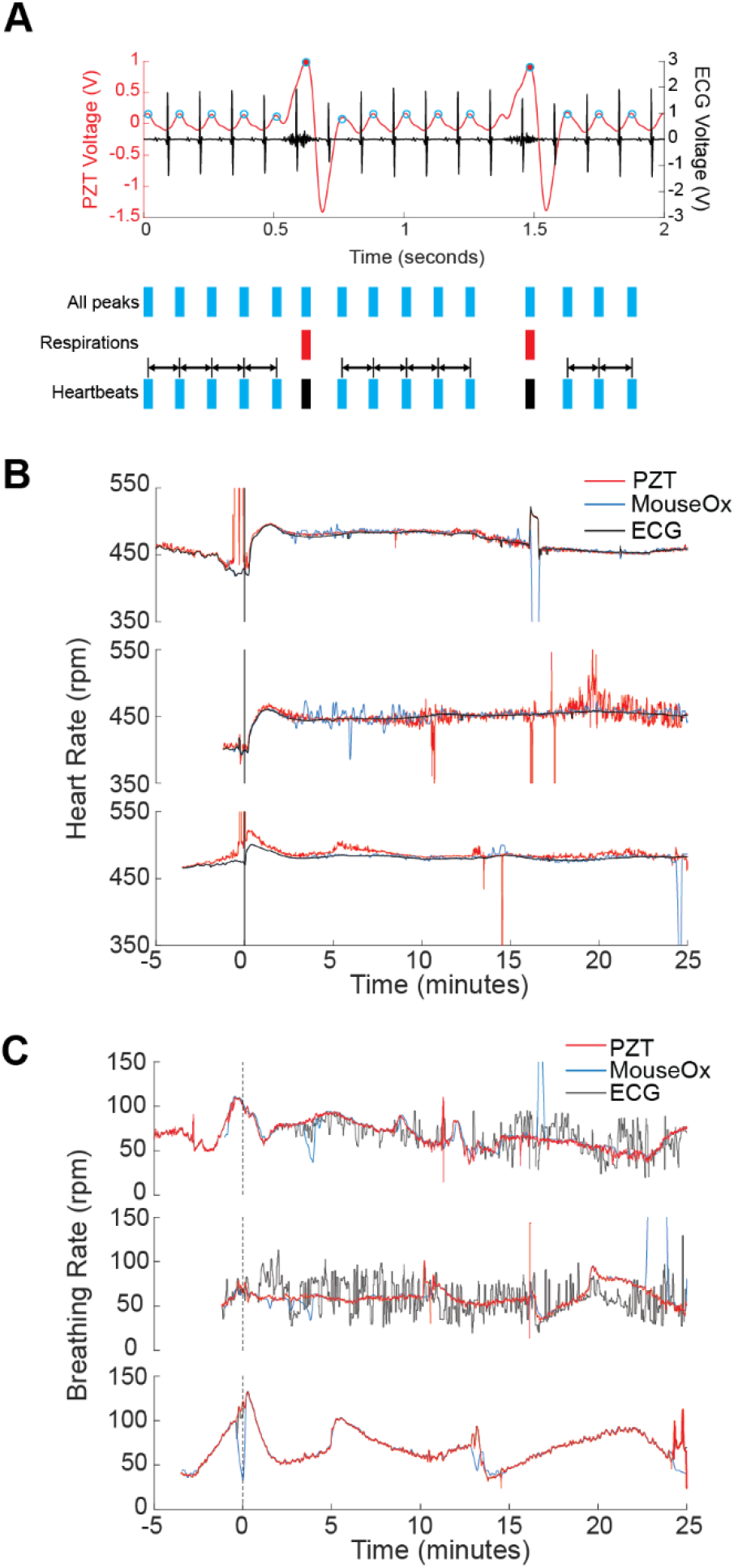
Detection of heart rate with MousePZT. **(A)** Simultaneous recording of animal ECG (black) and MousePZT signal (red) (top). The QRS complex precedes the peak in the MousePZT signal. Schematic of the method for extracting heart rate by detecting all peaks, cardiac and respiratory, then removing respiratory peaks, and determining the inter heart beat interval for the remaining cardiac peaks (bottom). **(B)** Heart rate changes in response to epinephrine (30-50 μL at 1 mg/mL, IM) for three different mice, as measured by MousePZT, ECG, and pulse-oximetry, simultaneously. All sensors show increases in heart rate following epinephrine administration. The MousePZT signal exhibits some motion artifact at the injection time in the first and third mice and a noisy heart rate measurement toward the end of the second recording. **(C)** Respiratory rate in response to epinephrine administration measured by MousePZT, ECG, and pulse-oximetry, simultaneously. The MousePZT and pulse-oximetry agreed across all three mice, while the ECG based respiratory rate quantification was noisy in two of the three mice.

For further confirmation of the heart rate detection, we administered epinephrine to isoflurane-anesthetized mice (n=3) to induce a predictable increase in heart rate while monitoring with the MousePZT, ECG, and a pulse-oximeter. All measurement approaches were broadly correlated, and clearly showed that heart rate increased after epinephrine administration. The heart rate measured using MousePZT was consistently a few bpm higher than that measured with ECG, likely due to the exclusion of heartbeat intervals during respiration and the use of a median, versus average, measurement. The overall relative error of PZT heart rate to ECG heart rate was 0.44 ± 0.8% and the overall absolute error was 2.0 ± 3.6 bpm. The pulse-oximeter relative and absolute errors relative to ECG were 0.20 ± 0.6% and 0.93 ± 2.7 bpm, respectively.

For respiratory rates measured using these three approaches, the signals matched well for one mouse, and for portions of the other two (Fig. 3C). The measurements taken using the MousePZT were smoother and less noisy compared to those from the pulse oximeter and ECG-derived signals, suggesting increased accuracy (Fig. 3C). These data show that the MousePZT system can provide a reliable measurement of median heart rate, in addition to measuring respiratory rate and providing respiratory trigger signals.

### Ketamine/Xylazine Monitoring for Heart Rate and Respiratory Rate

Finally, we assessed the performance of the MousePZT under ketamine/xylazine anesthesia and examined the relative value of measuring respiratory rate and heart rate (from ECG) in determining when an animal is waking up from anesthesia. We administered a single dose of a ketamine and xylazine solution to a mouse and tracked respiratory and cardiac rate following induction and until recovery, as determined by return of mobility. The mouse maintained a respiratory rate of 135 ± 8.5 respirations per minute (rpm) and a heart rate of 199 ± 7.9 bpm while anesthetized. Both heart and respiratory rate increased ~5 min before the mouse showed signs of ambulation. The respiratory rate, however, showed a detectible increase 51 s prior to the heart rate increase, and it increased at a faster initial rate of 2.2 rpm/s (1.6%/s), as compared to the heart rate increase of 1.5 bpm/s (0.76% /s) (Fig. 4). These results show that careful monitoring of respiratory rate can provide early and sensitive indicators of changes in anesthetic depth.

**Figure 4.**
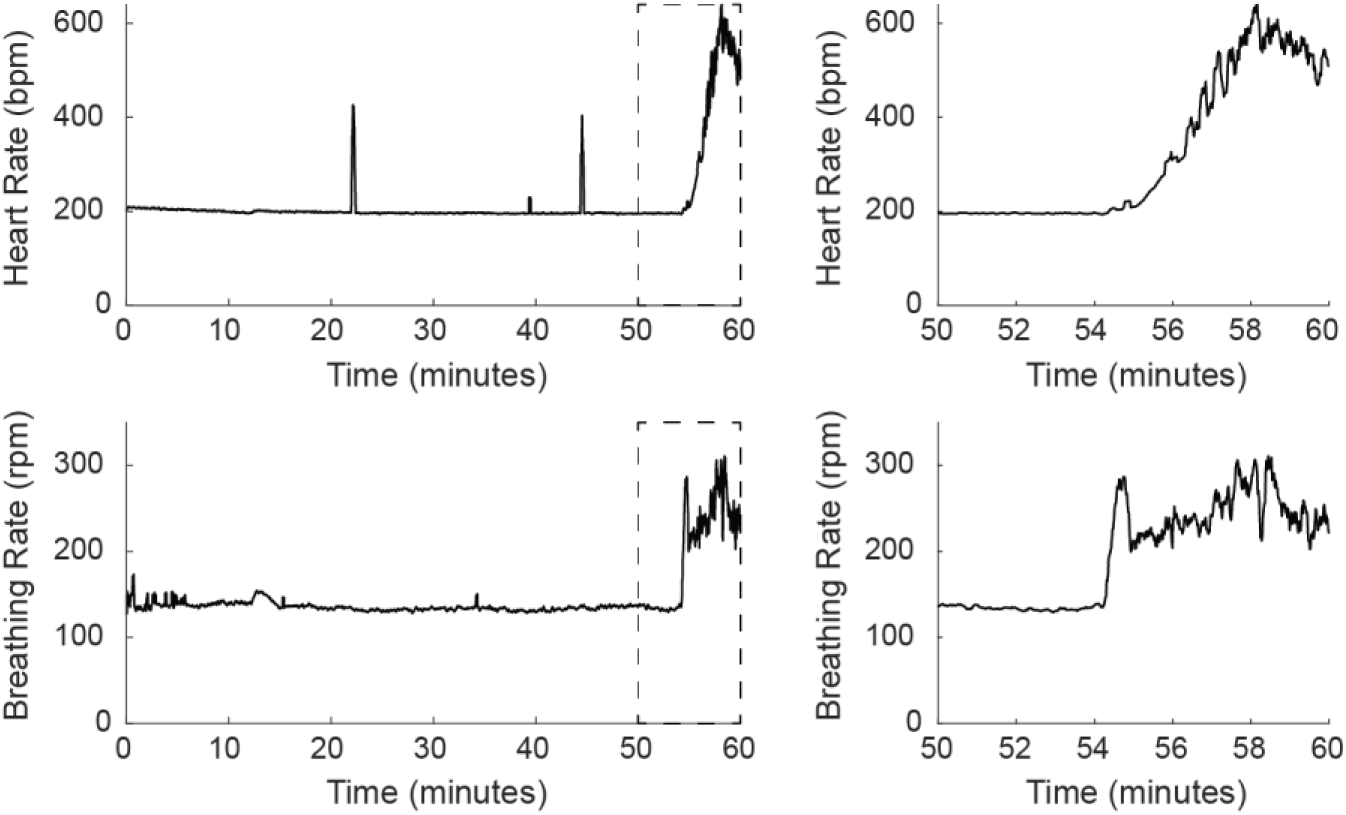
Respiratory rate changes provide an early indicator of anesthetic depth changes. Heart rate (top), as measured by ECG, and respiratory rate (bottom), as measured by MousePZT following anesthesia with ketamine/xylazine and continuing until ambulatory recovery at 60 min. Respiratory rate showed an earlier and faster increase than heart rate as seen in the final 10 min. prior to ambulation (right).

## Discussion

We have developed a system that allows for quantification of respiratory rate and heart rate in anesthetized mice using a simple sensor with additional circuitry to produce TTL pulses for respiratory gating that can be fabricated in-house at low cost. Signals can be obtained without need for electrode gel or insertion of electrode needles for ECG recordings, or hair removal for use of pulse-oximetry. The determined respiratory and heart rates closely followed those of commercial systems. In addition, the respiratory trigger signal was found to be highly accurate and required little tuning.

With our system, the MousePZT produced distinguishable and repeatable signal waveforms for respiratory rate measurements. Also using piezoelectric sensors the animal lays on, Gomes et al. noted that signals can be clearly identified with the animal in both ventral and dorsal recumbency (*18*). We observed similar signal clarity and additionally found a pronounced respiratory signal with the animal in lateral recumbency (Fig. 2A). For some animals and with some sensor placements, the waveform is less pronounced or differs in shape. We found common causes of reduced respiratory signal clarity to be due to the sensor being displaced over time, or from the animal laying on too soft of a base material. Adjustment of the animal or sensor position quickly restored the signal.

Other low-cost approaches for determination of respiratory rate have comparable reliability to our MousePZT system. The LED/photodiode combination used by Lemieux and Glover (*14*) gives very strong respiratory signals from the change in reflectivity due to the motion of the chest wall, providing both a means to measure respiratory rate and produce a respiratory trigger (although the implementation in Lemieux and Glover relied on a software comparator that could be improved). A thermistor implanted in the nose, used by McAfee et al. (*17*), produces more reliable signals than noninvasive temperature probes, but requires a surgical procedure. Without nasal epithelial implantation, thermistors can saturate following large breaths or sighs, leading to missed respirations (*23*).

While previous work has used piezoelectric signals to monitor respiratory rate, we add the capability for a respiratory trigger with MousePZT. The output TTL of the analog peak-detection circuit was robust and could be used for respiratory gating in applications such as CT or MRI to reduce respiratory motion artifacts. The sensitivity can be adjusted with a potentiometer to capture more or less of the respiratory waveform, detecting the start of inspiration or the transition from inspiration to expiration, respectively. The threshold was set manually for all experiments here but could be set adaptively by using a lowpass filtered (cutoff frequency < 0.1 Hz) modified Pan-Tompkins signal as the negative input to the comparator. While sensitivity can be increased to pick up some heartbeats, the hardware cannot pick up or distinguish between heartbeats and respirations. In addition, residual sensor noise may also be picked up intermittently with heartbeat detection. Therefore, in this paper, we focus on the use of the circuit for respiratory, but not cardiac, gating.

Heart beats also produce signals that can be picked up using a piezoelectric sensor, although they are more difficult to pick out than respiratory signals. Adding the capability to measure heart rate, without additional experimental or sensor complexity, provides another important vital sign to measure and record. With the sensor underneath the chest or abdomen and mouse in dorsal recumbency, clear peaks from heart beats could be detected. Our algorithm that determines the time between successive heart beats, while identifying and ignoring regions where respiration obscures heart beat signals, proved to provide a robust measurement of heart rate. Sato et al. (*23*) took a different signal processing approach to measure heart rate from piezoelectric sensors by applying a filter between 300 Hz and 1 kHz to capture the ‘heart sounds.’ We replicated this digitally but found it to be less reliable than our approach. ‘Heart sounds’ were often obscured by respiratory vibrations, or, at lower heart rates, the time between the heart valves closing to produce the sounds was increased, leading to irregular heartbeat detection. The MousePZT system matched the accuracy of more invasive ECG and pulse oximetry-based heart rate measurements, as long as at least two cardiac cycles occur in the interval between respiratory cycles. This occurs when the heart rate is at least 3-4 times higher than the respiratory rate. For this reason, heart rate was not measurable under ketamine/xylazine anesthesia, as heart rate (~200 bpm) was only 1.5 times higher than respiration rate (~135 rpm), which is typical for this anesthesia. Under isoflurane, heart rate is much faster, while respiration is similar or slower to ketamine/xylazine anesthesia, depending on anesthetic depth (*2*).

We showed that respiration rate is a strong indicator of anesthetic depth and found it increased sooner and faster than heart rate in an animal waking up from ketamine/xylazine anesthesia. Other studies have similarly demonstrated that respiration rate is heavily influenced by anesthetic depth (*1, 2, 10, 18*), making this vital sign a robust metric to monitor.

In conclusion, we have demonstrated that a simple, low-cost sensor can be utilized for monitoring multiple vital signs in anesthetized mice with high accuracy across a range of sensor positions, animal orientations, and anesthetics used. MousePZT more accurately measured respiratory rate and performed as well in determining the heart rate as compared to other, more complex, monitoring methods, such as ECG and pulse-oximetry. Through the application of a simple peak-detection circuit, we further demonstrated high sensitivity and specificity output of a respiratory-locked trigger signal. MousePZT is an effective tools for of vital sign monitoring in anesthetized mice and is particularly useful when experimental conditions (e.g., imaging under low light) make direct visualization of the animal difficult. All resources to enable construction of a MousePZT, including a detailed parts list, a step-by-step construction guide for assembly of the circuit and overall system, ready-to-use design files for the printed circuit boards and the 3D printed housing, all microcontroller code and computer software, as well as a troubleshooting and debugging guide, are available at github.com/sn-lab/Mouse-Breathing-Sensor.

## Materials and Methods

### MousePZT system

#### Sensor Circuit

An Arduino microcontroller serves as the data acquisition board and 5 V power supply for the circuit. A virtual ground is created with two 10-kOhm resistors and a single-supply op-amp as a buffer. The sensor is a 27-mm diameter piezoceramic bender (PUI Audio, Inc.) with two soldered leads and electrical tape as insulation. The PZT is tied to the virtual ground via a 1-MOhm resistor. The signal from the PZT is amplified through a noninverting amplifier (gain = 3 [9.5dB]) before passing through a secondary inverting amplifier and low-pass filter (gain = 2 [6dB], fc = 100 Hz), followed by an adjustable twin-t notch filter centered on the line noise frequency (on-board potentiometers tune this for 50-Hz or 60-Hz line noise). The signal is finally passed through a noninverting amplifier with a variable gain set by a potentiometer and read out by the on-board analog-to-digital converter of the microcontroller. A simplified experimental setup and schematic of the MousePZT circuit are depicted in Fig. 1A and B, respectively.

#### PZT-Derived Respiratory Rate and Heart Rate Calculation

The microcontroller communicates with an app written in MATLAB’s App Designer via a serial port and transmits the signal voltage at a sampling rate specified by the app. The baud rate is set to 115,200 bits/s to balance speed and stability, limiting the max theoretical sampling frequency (fs) due to communication to approximately 7.2 kHz. As the signals of interest are low frequency (HR < 10 Hz), the signal data is boxcar filtered with the window size (ws) set to remove the line noise and resulting harmonics completely (ws = fs/[line frequency]). Once per second, MATLAB takes a 5-s window of the boxcar-filtered data and locates the respiratory peaks using findpeaks(). Detected respiratory peak intervals are averaged to obtain the respiration rate for that 5-s window. For determination of heart rate from the PZT signal, respiratory and heart rate peaks are detected from the digitally filtered signal without additional processing using findpeaks(), with the peak prominence and minimal distance between peaks settings significantly reduced to enable detection of both heart beat associated peaks and respiratory peaks. Detected peak locations that overlap with the initial respiratory peaks are converted to NaNs as the respiratory signal dominates the heart signal in those intervals. The heart rate is then calculated from the median of the time interval between successive heart beats, with NaN intervals omitted (Fig. 3A).

#### Analog Respiratory Trigger Circuit

For analog detection of respiratory peaks, a modified Pan-Tompkins analog circuit (*24*) (Fig. 1C) was implemented in which the signal is differentiated, rectified (not squared), and lowpass filtered. To avoid saturation, the output respiratory signal of the sensor circuit is reduced to 1/18 of the original amplitude using a voltage divider to the virtual ground and buffered before being differentiated with corner frequencies of 16 Hz and 33 Hz and a resulting gain of 44 [33 dB]. The differentiated signal is sent through a full-wave rectifier before being low passed at 8.7 Hz. A rectifier was opted for rather than a squarer circuit to reduce circuit complexity and avoid exceeding the voltage range available. A comparator outputs a TTL pulse when the processed signal exceeds a reference voltage set by a potentiometer. The voltage signals at each stage of this circuit are depicted in Fig. 1D. Before each recording with the analog peak detection, the two potentiometers in the circuit were used to adjust the signal amplitude and trigger sensitivity. The potentiometer for the output signal was adjusted to produce a 1-2 V peak-to-peak amplitude due to respiration, as viewed in the Matlab app or on an oscilloscope. The second potentiometer for the respiratory peak-detection circuit was adjusted to avoid erratic triggering (an on-board light emitting diode (LED) flashes with each trigger to facilitate easy tuning) and to either lower the threshold to enable triggering across the entire respiratory waveform or increasing the threshold to trigger just at the transition from inspiration to expiration.

### In vivo validation studies

#### Animals

All animal procedures were approved by the Cornell Institutional Animal Care and Use Committee (protocol 2015-0029) and were performed under the guidance of the Cornell Center for Animal Resources and Education. We used seven adult C57BL/6 mice of both sexes between 3 and 8 months of age. All mice recovered from the procedures performed in this study.

#### Isoflurane induction and adjustment

Anesthesia was induced with 3% isoflurane in medical air using an induction chamber. Once unresponsive to toe or tail pinch, the animal was moved to a stereotactic frame with a nose cone and a feedback-controlled heating pad (40-90-8D, FHC) set to 37°C. Isoflurane was reduced to 1-2% for maintenance and adjusted to vary the respiratory rate during recordings. The respiratory rates of the mice were visually and digitally monitored by the experimenter and with the MousePZT system, respectively.

#### Setting and verifying the respiratory peak detector

To determine the accuracy of the respiratory trigger, the output PZT signal and peak detector output TTL signals were recorded for 60-s epochs on a USB-6003 DAQ board (National Instruments). The modified Pan-Tompkins algorithm was applied to the respiratory signal and the respiratory peaks and peak widths were determined using findpeaks() in MATLAB. The signal was split into respiratory and inter-respiratory epochs based on the respiratory peak locations and respective widths. Each epoch was then examined for high or low TTL values to determine the true positive (TTL during respiration), true negative (no TTL outside respiration), false positive (TTL outside respiration), and false negative (no TTL during respiration) rates. For calculation of respiratory rate using the trigger signal, we applied a triangular moving average filter on the TTL signals before differentiating and running the findpeaks() function to identify the rising pulse edges for calculation of the TTL-based respiratory rate. Alternatively, for visual accuracy of the peak-detector, a LED was tied to the output of the comparator to indicate when the output trigger was high, and the mouse and LED were recorded at ~30 fps using a cell phone camera. The video was stabilized in FIJI (ImageJ) using the Linear Alignment with SIFT plugin and cropped so that reviewers could score the start and stop of respiratory cycles without input bias from the LED. A region-of-interest (ROI) was placed over the LED in the stabilized video to determine, based on pixel intensity, when the trigger was active.

#### Comparative heart rate and respiratory rate monitoring after pharmacological manipulation of heart rate

Three electrode needles were subcutaneously attached to the left forelimb, right forelimb, and left hindlimb of isoflurane-anesthetized mice. ECG traces were collected from Lead I, based on the Einthoven’s triangle configuration. The leads were fed into a bioamplifier (ISO-80, World Precision Instruments) with gain set to 10^3^ [60 dB] and the bandpass filter set to 0.1-300 Hz. Two mice were recorded on a USB-6003 DAQ board without additional intervention. Three mice were recorded while also incorporating a thigh-placed pulse-oximeter (MouseOx, StarrLife Sciences) to track heart rate and respiration rate. In preparation for use of the pulse-oximeter, hair was removed from the left hind limb using a small animal shaver and depilatory cream (Nair). MousePZT and ECG signals were recorded at 10 kHz on the DAQ board and the calculated pulse oximeter vital signs at 15 Hz using MouseOx software. DAQ-recorded signals were synced with the MouseOx output by running a cross-correlation on the rising edge of the heart rate between the ECG-derived rates and MouseOx-derived rates. Mice were recorded for up to 5 minutes to achieve a baseline rate, followed by intramuscular administration of 30-50 μL epinephrine (1 mg/mL) into the right hindlimb to induce a predictable increase in heart rate. Mice were recorded for at least 30 minutes following epinephrine injection.

#### ECG-Derived Heart and Respiratory Rate Calculation

Heart rate was determined by detection of the QRS complexes in the ECG signal (*25*). Typically, these peaks have a large single peak and therefore can be detected simply using MATLAB’s findpeaks() function. The peak-to-peak interval is then averaged across 5-s windows to produce a heart rate measurement that is compared to that from the PZT sensor. In cases where the J wave nearly matches or exceeds the R wave amplitude, resulting in erroneous heart rate determination (*25*), we apply the Pan-Tompkins algorithm with corner frequencies at 30 and 100 Hz, and a boxcar filter of 10 ms for the squared derivative. ECG signals have been used to reproduce respiratory waveforms and to calculate respiration rate using the fluctuations in the R-R interval (heart rate increases slightly during inspiration), or by examination of the R-peak amplitude (decreases during inspiration) over time (*26*). While we examined both the R-R interval and R-peak amplitudes, we found the R-peak amplitude to be more reliable due to the higher sampling rate required for R-R interval measurements in mice due to the short interval between successive heart beats. We apply a Modified-Akima 1D interpolation to the R-peak amplitudes before detrending to generate a signal that varies with respiration, from which we determine respiratory rate over a 5-s window.

#### Monitoring under Ketamine/Xylazine Administration

One mouse was given a single dose of ketamine/xylazine solution (0.1 mL/10 g body weight; 10 mg/mL ketamine and 1 mg/mL xylazine) intraperitoneally. Once unresponsive to toe or tail pinch, the mouse was placed in ventral recumbency on a heating pad with temperature feedback set to 37°C. The PZT sensor was then placed between the thorax and the heating pad. For ECG recording, to avoid damaging the skin or causing discomfort to the animal upon recovery, electrode needles were placed in light contact with the left and right forepaws and the left hindpaw with electrode gel for signal transduction. ECG and PZT signals were recorded on the USB-6003 DAQ board at 10 kHz until the animal showed signs of ambulation, about 75 minutes after the injection.

## Supporting information

Recording1

## Acknowledgements

We thank Jordan Harrod, Riona Reeves, Rohan Roy, Julia Telischi, and Kelly Wilson for their early contributions in the design of this device.

## Author contributions

Conceptualization: DAR, CBS

System Design: DAR

Investigation: DAR, AEB

Software: DAR

Formal analysis: DAR, AEB

Funding acquisition: CBS

Writing – original draft: DAR

Writing – review & editing: DAR, AEB, CBS

## Data and materials availability

The data underlying each figure or result will be uploaded to Cornell’s eCommons and will be publicly available. Software and PCD design will be uploaded to github.com/sn-lab/

